# PSMC3 is required for spermatogonia niche establishment in mouse spermatogenesis

**DOI:** 10.1101/529149

**Authors:** Maria Cristina S. Pranchevicius, Luciana Previato, Rodrigo O. de Castro, Roberto J. Pezza

## Abstract

Males produce millions of spermatozoa each day, which are originated from spermatogonia. Spermatogonia niche establishment and maintenance and the subsequent haploidization of spermatocytes in meiosis are hallmarks of this process. The function of the individual players and coordinated mechanisms regulating different stages of gametogenesis in mammals are not well understood. In this work we focused on the role of PSMC3 in mouse gametogenesis. We observed that *Psmc3* is highly expressed in mouse testis, and it is widely expressed in different stages of gamete formation. Conditional deletion of *Psmc3* results in both male and female impairment of gonad development at early pre-meiotic stages, but has no apparent effect on meiosis progression. This is likely a consequence of abnormal spermatogonia niche establishment and/or maintenance, revealed by a massive loss of undifferentiated spermatogonia. Our work defines a fundamental role of PSMC3 functions in spermatogenesis during spermatogonia development with direct implications in fertility.

## Introduction

Spermatogenesis is the process by which diploid spermatogonia cells produce haploid mature gametes called spermatozoa. During this process, spermatogonia niche establishment and maintenance are essential to ensure that the genetic information is passed to the next generation. Indeed, errors in producing or differentiating spermatogonia can result in male infertility. Spermatogenesis begins with primordial germ cells migration to the developing male gonad and division to produce an undifferentiated stem cell type of spermatogonia (type A). The latter cells divide and some of them generate differentiated type B spermatogonia, the last to undergo mitotic division and the precursor of primary spermatocytes. Primary spermatocytes undergo meiosis to half their chromosome complement and yield a pair of secondary spermatocytes, which differentiate to produce haploid gametes [1].

In order to explain possible causes of infertility, we need to identify and understand the function of the individual players and coordinated mechanisms regulating mammalian spermatogenesis. Here, we focus on understanding the role of PSMC3 in mouse germ cell development. Analysis of mouse PSMC3 (a.k.a. TBP1/Tat-binding protein 1, RPT-5 or S6A) protein sequence indicates that PSMC3 belongs to the AAA ATPase family of proteins. Starting from the amino-terminus, this protein features a putative leucine zipper motif with possible DNA binding activity; an ATPase Walker A motif, an ATPase Walker B motif; and a putative helicase domain with a DEXD motif, that relates PSMC3 to the superfamily 2 DEAH helicases.

PSMC3 has been associated with a number of different cell functions. This includes participating in the 19S regulatory subunit of the proteasome [2], whose main function is degradation of excess, no longer needed, and defective proteins. In most organisms proteasome activity acts via the ubiquitin/26S proteasome system [3, 4]. This system involves the specific attachment of a chain of ubiquitin to the protein target by the E1-E3 enzymes [3]. The 26S proteasome complex recognizes labeled target proteins. This complex can be subdivided into a 20S core protease and a 19S regulatory part [5]. The 19S subunit works by recognizing the target proteins and delivering them to the 20S subunit for degradation. Among other proteins the 19S subunit is composed by the AAA-ATPases Psmc 1-5 (Rpt1-Rpt6) (regulatory particle triple A ATPase) [6]. All Psmc/Rpt proteins are essential in yeast [7] and they form an hexameric ATPase complex [8, 9]. In *Arabidopsis*, mutants affecting 19S RP ATPase subunits show severe defects in maintaining the pool of stem cells in the root. Importantly, gametophyte development requires proteasome function, which is evident by chemical inhibition of the proteasome resulting in pollen developmental defects [10–12]. Recent studies using insertion mutants affecting proteasome components observed that alleles affecting *Rpt5a* (an *Arabidopsis* ortholog of *Psmc3*) displayed severe male gametophyte development defects, with pollen development arrested before cells enter meiosis at the second pollen mitosis stage [13].

PSMC3 has also being implicated in different cellular events that do not require proteolysis such as transcriptional initiation and elongation [14–16], DNA repair [17], and as a negative regulator of cell proliferation [18, 19]. By comparing gene expression profiles from normal and abnormal human testes with those from comparable infertile mouse models, a number of genes critical for male fertility have been identified [20]. Among the expression of 19 human genes that were different between normal and abnormal samples, *Psmc3* appears as a top candidate [20]. Another lead to the function of PSMC3 independent of the proteasome is that the HOP2/TBPIP protein, a strong interaction partner of PSMC3, has an extensively documented role in proper meiotic chromosome segregation and fertility through its interaction with DMC1 and RAD51, both central components of the recombination pathway [21–24].

The role of PSMC3 in mammalian spermatogenesis has not been explored. Herein, we present the phenotype associated to the conditional deletion of *Psmc3* in mouse gonads. Males are infertile likely a result of absence of any gametocyte type in the gonad. Meiosis is apparently not affected in *Psmc3^-/-^* mice when ablation of PSMC3 occurs at meiotic stages. However, testis development is impaired at early stages during spermatogonia niche establishment, with massive loss of undifferentiated spermatogonia. Our work in understanding the functions of PSMC3 in germ cell development has broader implications in defining mechanisms responsible for infertility.

## Materials and Methods

### Mice and Genotyping

Experiments involving mice conformed to relevant regulatory standards and were approved by the IACUC (Institutional Animal Care and Use Committee).

Mice: The *Psmc3* stem cells carrying a floxed allele mice was obtained from International Knockout Mouse Consortium. Transgenic *Cre* recombinase mice *Ddx4-Cre*^FVB-Tg(Ddx4-cre)1Dcas/J^ or *Stra8-iCre*^B6.FVB-Tg(Stra8-icre)1Reb/LguJ^ were purchased from The Jackson Laboratory (Bar Harbor, ME). *Spo11*-Cre mice were provided by Dr. P. Jordan (Johns Hopkins University Bloomberg School of Public Health, Baltimore, MD). All mice were maintained on a mixed genetic background at the Laboratory Animal Resource Center of Oklahoma Medical Research Foundation. All animal work was carried out in accordance with IACUC protocols.

Genotyping: characterization of wild type and floxed alleles was carried out by PCR using the following oligonucleotides (see Fig. 2A): 1F 5’- CAAGCAGATCCAGGAGGTAAG, 1R 5’- CATGGCTCAGAGAGTAAGAGTG, 2F 5’- CATGTCTGGATCCGGGGGTA, 2R 5’- CCTACTGCGACTATAGAGATATC, 3F 5’- GGATTCCAGAGAGATTGGAGATTGT, 3R 5’- CCTACTGCGACTATAGAGATATC, 4F 5’- GGATTCCAGAGAGATTGGAGATTGT, and 4R 5’- GAACGGGCCACACAAATCTAGTA. The presence of *cre* recombinase allele was determine by PCR using the following primers: *Spo11*-Cre forward 5’-CCATCTGCCACCAGCCAG, *Spo11*-Cre reverse 5’-TCGCCATCTTCCAGCAGG, *Ddx4-*Cre forward 5’- CACGTGCAGCCGTTTAAGCCGCGT, *Ddx4*-Cre reverse 5’- TTCCCATTCTAAACAACACCCTGAA, *Stra8*-Cre forward 5’- AGATGCCAGGACATCAGGAACCTG and *Str8*-Cre reverse 5’- ATCAGCCACACCAGACAGAGAGATC.

**Figure 2.**
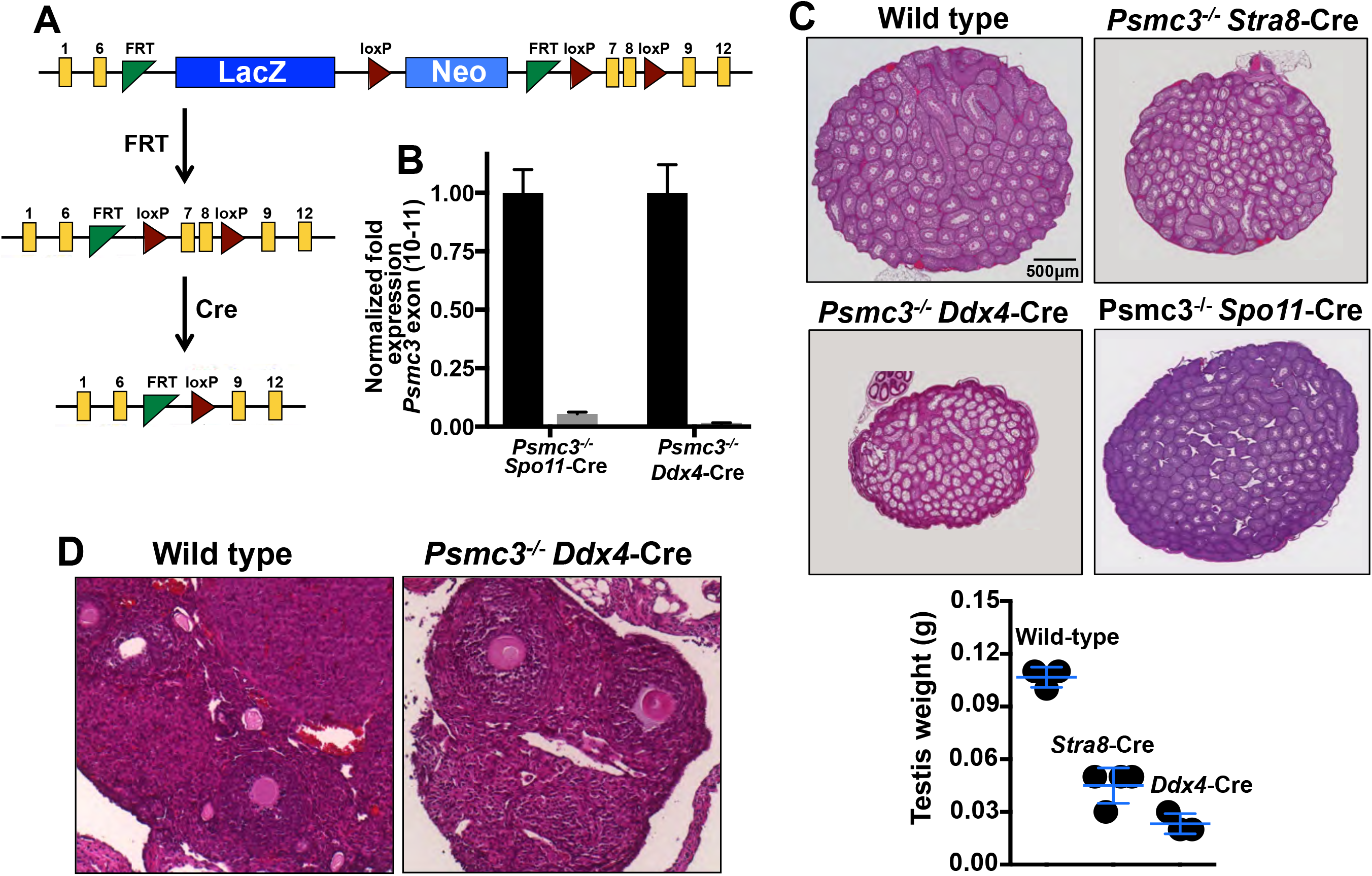
*Psmc3* gene targeting design and testis size phenotype of *Psmc3* mutant mice. Testis specific Cre knockout strategy for deletion of *Psmc3*. A trapping cassette was inserted to delete exons 7 and 8 in Psmc3. **(B)** *Psmc3* transcription levels expression in whole testis of wild type and mutant (*Psmc3^-/-^, Spo11*-Cre) mice and enriched fractions of spermatogonia cells from wild type and mutant (*Psmc3^-/-^*, *Ddx4*-Cre) mice assessed by RT-PCR. **(C)** PLZF&E stained testis of wild type, *Stra8*-*Psmc3^−/−^*, *Ddx4*-*Psmc3^−/−^,* and *Spo11*-*Psmc3^−/−^* mice. Quantification of testis weight for wild type and homozygous knockout mice is also shown. **(D)** PLZF&E stained ovaries of wild type and *Ddx4*-*Psmc3^−/−^* mice.

### Histology and immunostaining

Testes and ovaries were dissected, fixed in 4% paraformaldehyde and processed for paraffin embedding. After sectioning (5–8-μm), tissues were positioned on microscope slides and analyzed using hematoxylin and eosin using standard protocols. For immunostaining analysis, tissue sections were deparaffinized, rehydrated and antigen was recovered in sodium citrate buffer (10 mM Sodium citrate, 0.05% Tween 20, pH 6.0) by heat/pressure-induced epitope retrieval. Incubations with primary antibodies were carried out for 2 PLZF at 37°C in PBS/BSA 3%. Primary antibodies used in this study were as follows: monoclonal mouse antibody raised against mouse SOX9 at 1∶:500 dilution (AbCam, ab26414), polyclonal rabbit antibody raised against mouse STRA8 at 1∶:500 dilution (AbCam, ab49602), polyclonal rabbit antibody raised against mouse TRA98 at 1∶:200 dilution (AbCam, 82527), monoclonal mouse antibody raised against mouse PLZF at 1∶:50 dilution (Santa Cruz, 28319). Following three washes in 1× PBS, slides were incubated for 1 PLZF at room temperature with secondary antibodies. A combination of Fluorescein isothiocyanate (FITC)-conjugated goat anti-rabbit IgG (Jackson laboratories) with Rhodamine-conjugated goat anti-mouse IgG and Cy5-conjugated goat anti-human IgG each diluted 1∶:450 were used for simultaneous triple immunolabeling. For Stra8, we used the ImmPRESS™ Reagent Anti-Rabbit IgG Peroxidase (Vector Laboratories) and hematoxylin as counterstaining. Slides were subsequently counterstained for 3 min with 2µg/ml DAPI containing Vectashield mounting solution (Vector Laboratories) and sealed with nail varnish.

### Statistical Analyses

Results are presented as mean ± standard deviation (SD). Statistical analysis was performed using Prism Graph statistical software. Two-tailed unpaired Student’s *t*-test was used for comparisons between 2 groups. P< 0.05 was considered statistically significant.

## Results

### *Psmc3* expression in mouse testis

Tissue specific expression and kinetics of expression in testis may help reveal the function/s of PSMC3. We performed RT-PCR on total purified RNA from different mouse tissues and specific primers designed to analyze the level of expression of *Psmc3* (Figure 1A). Our results are consistent with previous reports [25] and show that *Psmc3* is highly expressed in testis and a relative minor amount in other tissues such as thymus, brain, liver, and kidney.

**Figure 1.**
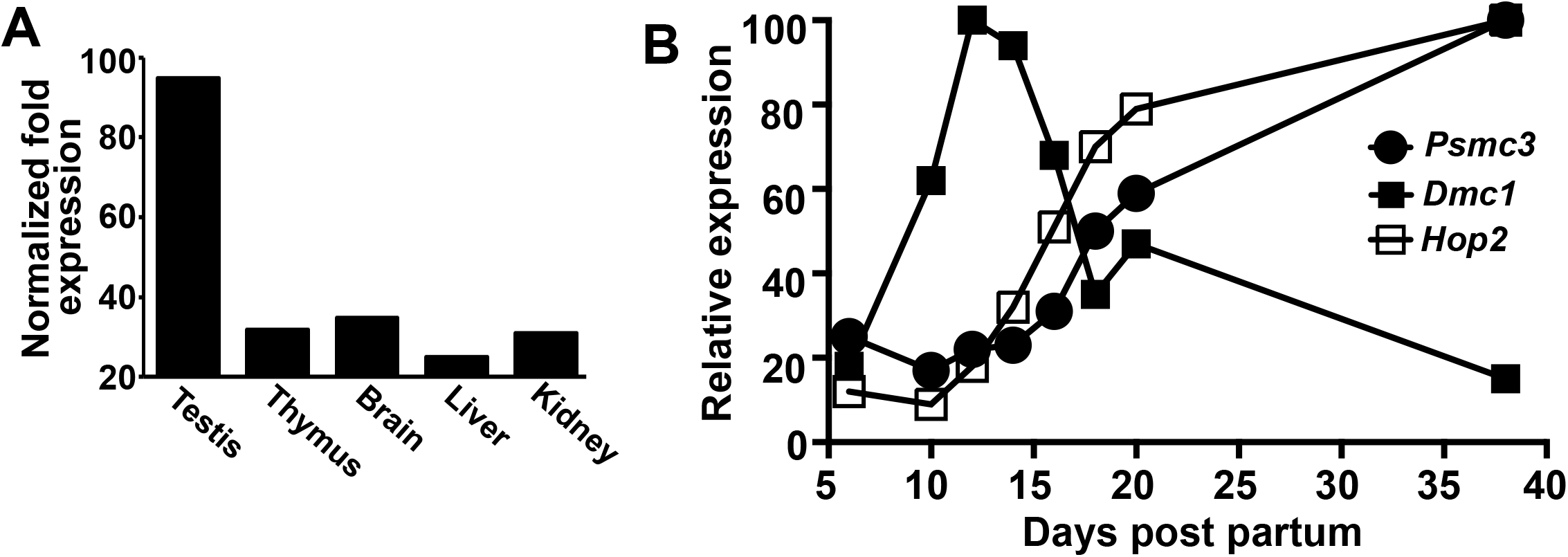
*Psmc3* expression during gametogenesis. **(A)** Expression of *Psmc3* in different tissues assessed by RT-PCR **(B)** Kinetics of *Psmc3* gene expression in testis of 6, 10, 12, 14, 16, 18, 20, and 38 dpp mice. Expression of *Psmc3* and *Dmc1* and *Hop2* was assessed by RNA-seq.

We then used a complete data set of gene expression previously generated in testis [26] to determine *Psmc3* expression during testis development (Figure 1B). We used *Dmc1* and *Hop2* expression times as markers for proteins expressed during early meiotic specific stages of gamete development. We concluded that *Psmc3* is initially expressed at pre-meiotic stages of gamete development (6 days post partum (dpp) samples) and gradually increases after 10 dpp as spermatogenesis progresses, which may suggest a dual role for *Psmc3* both at early and later stages of gamete development.

### Generation of *Psmc3* testis specific knockout mice

We generated *Psmc3* knockout mice using germline conditional inactivation. Two FRT sites are located in between exons 6 and 7 and flank LacZ, a loxP site, and a Neo cassette. Exons 7 and 8 of *Psmc3* are flanked by two loxP sites (Fig. 2A). Heterozygous mice carrying *Psmc3* FRT sites and floxed 7 and 8 alleles were first mated with transgenic mice expressing FRT recombinase. Products of this cross were mated with *Ddx4*-, *Stra8*-, or *Spo11*-Cre transgenic mice (Fig. 2B). Cre activity in *Ddx4*-Cre mice first becomes detectable in primordial germ cells (embryonic day 15.5). Thus, conditional knockout mice *Ddx4*-Cre; *Psmc3^fl/Δ^* (here called *Ddx4*-*Psmc3*^−/−^) was generated by crossing males *Ddx4*-Cre; *Psmc3^WT/Δ^* with females *Psmc3^fl/fl^* mice. The *Stra8*-Cre is expressed later in differentiated spermatogonia cells, which allows the deletion of the floxed allele only in cells already committed to meiosis (*Stra8*-Cre; *Psmc3^fl/Δ^*, *Stra8-Psmc3^−/−^*). Finally, we generated mutants using *Spo11*-Cre (*Psmc3^fl/Δ^*, *Spo11-Psmc3*^−/−^) mice in which the floxed *Psmc3* allele is expected to be deleted only in early primary spermatocytes. We confirmed deletion of *Psmc3* by RT-qPCR (Fig. 2B).

### Deletion of *Psmc3* results in testis developmental defects

If *Psmc3* participates in any stage of gamete development, we expect that their deletion will lead to disruption of gametogenesis, which can be studied by comparative tissue analysis of wild type and mutant testis. *Psmc3^−/−^* adult mice appear normal in all aspects except in reproductive tissues (Fig. 2C). However, *Stra8-*Psmc3^−/−^ (0.045g ± 0.005, n = 4, P=0.0002, t test) and *Ddx4*-*Psmc3^−/−^* males (0.023g ± 0.003, n = 3, P≤0.0001, t test) had significantly smaller testes than wild type (0.11 g ± 0.003, n = 3) littermates, with *Ddx4*-*Psmc3^−/−^* showing the most significant reduction in testis *size* (Fig. 2C). This substantial reduction in size is an indication of severe developmental defects in testis. In contrast, *Spo11*-*Psmc3^−/−^* (0.1 g ± 0.0063, n = 4, P=0.1, t test) did not show any significant difference compared to wild type littermates (Fig. 2C). *Spo11* is expressed in early prophase I, during leptonema [27]. Therefore, normal testis size in *Psmc3^−/−^ with Spo11*-Cre background indicates that developmental defects triggered by PSMC3 depletion are originated in pre-meiotic stages of gamete development and that PSMC3 has no apparent role during mouse meiosis. Thus, homozygous *Stra8*- and *Ddx4*-*Psmc3^−/−^* mutant mice show severe blocks of spermatogenesis. We also analyzed hematoxylin-eosin histological sections of 45-days-old wild type and *Ddx4*-*Psmc3^−/−^* ovaries. We note that albeit a significant reduction occurred in ovary size and increase in stromal cells, a reduced number of follicles can be observed in the *Ddx4*-*Psmc3^−/−^* mice (Fig. 2D). We conclude that Psmc3 plays a role in male and female gametogenesis.

Detailed tissue analysis indicates that both *Stra8*- and *Ddx4*-*Psmc3^−/−^* males develop testicular hypoplasia with hyperplasia of interstitial cells and a lack of spermatozoa, with *Ddx4*-*Psmc3^−/−^* showing the most severe phenotype (Fig. 3A and B). Although there were no alterations in number of seminiferous tubules, the diameter of seminiferous tubules was reduced (wild type, 319.8μm ± 13.26, n=30; mutant, 173.6 μm ± 21.87, n = 30). Spermatids represent the most advanced spermatogenic cells in the *Stra8*-*Psmc3^−/−^* mice, indicating that spermatogenesis progress, albeit with a severely reduced number of cells (wild type average 53 cells per seminiferous *tubules* while *Stra8-Psmc3^−/−^* average 32 cells per seminiferous tubule). Analysis of *Ddx4*-*Psmc3^−/−^* revealed near total loss of germ cells in seminiferous tubules. Although no meiocytes were observed, even those cell types at early development (i.e. spermatogonia); Sertoli cells were apparently not affected (Fig. 3A and B).

**Figure 3.**
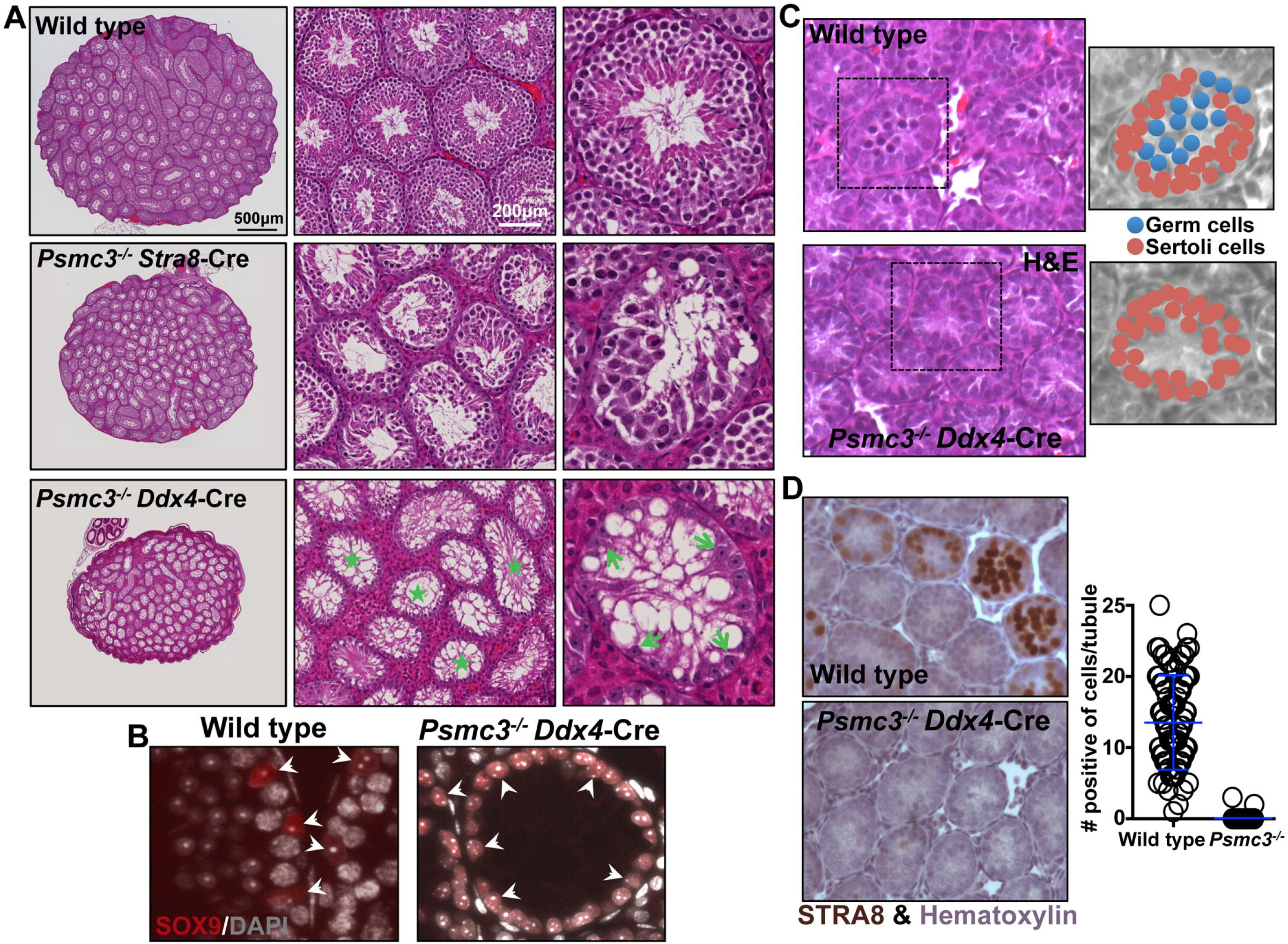
*Ddx4*-*Psmc3^−/−^* mice show profound defects in gametogenesis. **(A)** Details of histological sections stained with PLZF&E of wild type, *Stra8-Psmc3^−/−^*, and *Ddx4-Psmc3^−/−^* seminiferous tubules. Stars mark seminiferous tubules with absent germ cells. Note unchanged number and morphology of Sertoli cells (indicated by green arrows). **(B)** Histological sections of wild type and *Ddx4-Psmc3^−/−^* testis immunostained with SOX9 (to mark Sertoli cells) and DAPI (to mark nuclei). **(C)** PLZF&E stained 9dpp testis of wild type and *Ddx4*-*Psmc3^−/−^* mice. The inserts show magnification of seminiferous tubules and composition and distribution of cells. **(D)** STRA8 immunostained and hematoxylin stained 9dpp testis of wild type and *Ddx4*-*Psmc3^−/−^* mice. Starts indicate positive seminiferous tubules. Quantitation of number of cells STRA8 positive per positive seminiferous tubule is also shown.

### PSMC3 is apparently required for spermatogonia niche establishment and maintenance

The severe phenotype observed in *Ddx4-Psmc3^−/−^* mice (Fig. 2 and 3A and B) prompted us to explore earlier stages of testis development with the premise that morphological changes between the mutant and wild type may reveal differences in early spermatogenesis differentiation. Testis from mice at different postnatal ages were collected, paraffin embedded, and tissue slides analyzed by PLZF&E or immunohistochemistry. PLZF&E stained testis sections from 9dpp *Ddx4*-*Psmc3*^−/−^ show near absence of gametocyte and differences in cell composition compared to wild type (Fig. 3C). To analyze this in detail, we immunostained testis sections with Stra8, which marks differentiating spermatogonia (Fig. 3D). While several tubules in wild type contain cells expressing STRA8 (12.83 average number of cells per positive tubule, n=34 seminiferous tubules), tubules in *Psmc3*^−/−^ samples show near absence of positive cells for these markers (0 STRA8 positive cells, n=50) (Fig. 3D). These results suggested that testis developmental defects in *Psmc3*^−/−^ mice begin early during pre-meiotic stages of postnatal development, possibly before spermatogonia differentiate, and are the cause of absent germ cells in adult mutant mice.

To investigate this in detail, we then analyzed 1, 3, 5, 7 and 9 dpp testis sections by immunostaining with antibodies specific for PLZF/ZBTB16, used as a marker for undifferentiated spermatogonia, and TRA98, which mark all germ cells (Fig. 4A). Similar number of cells are positive for both markers as observed for both wild type and *Psmc3* mutant in 1dpp (PLZF wild type, average ± standard deviation, 1.0 ± 1.3, n = 123 seminiferous tubules; mutant, 1.6 ± 1.7, n = 73, P=0.014, t test) (TRA98 wild type, 0.33 ± 0.73, n = 93; mutant, 0.08 ± 0.3, n = 73, P=0.007, t test) and 3dpp (PLZF wild type, 2.0 ± 1.3, n = 66 seminiferous tubules; mutant, 2.0 ± 1.6, n = 23, P=0.89, t test) (TRA98 wild type, 0.94 ± 0.97, n = 66; mutant, 1.4 ± 1.02, n = 23, P=0.05, t test) testis.

**Figure 4.**
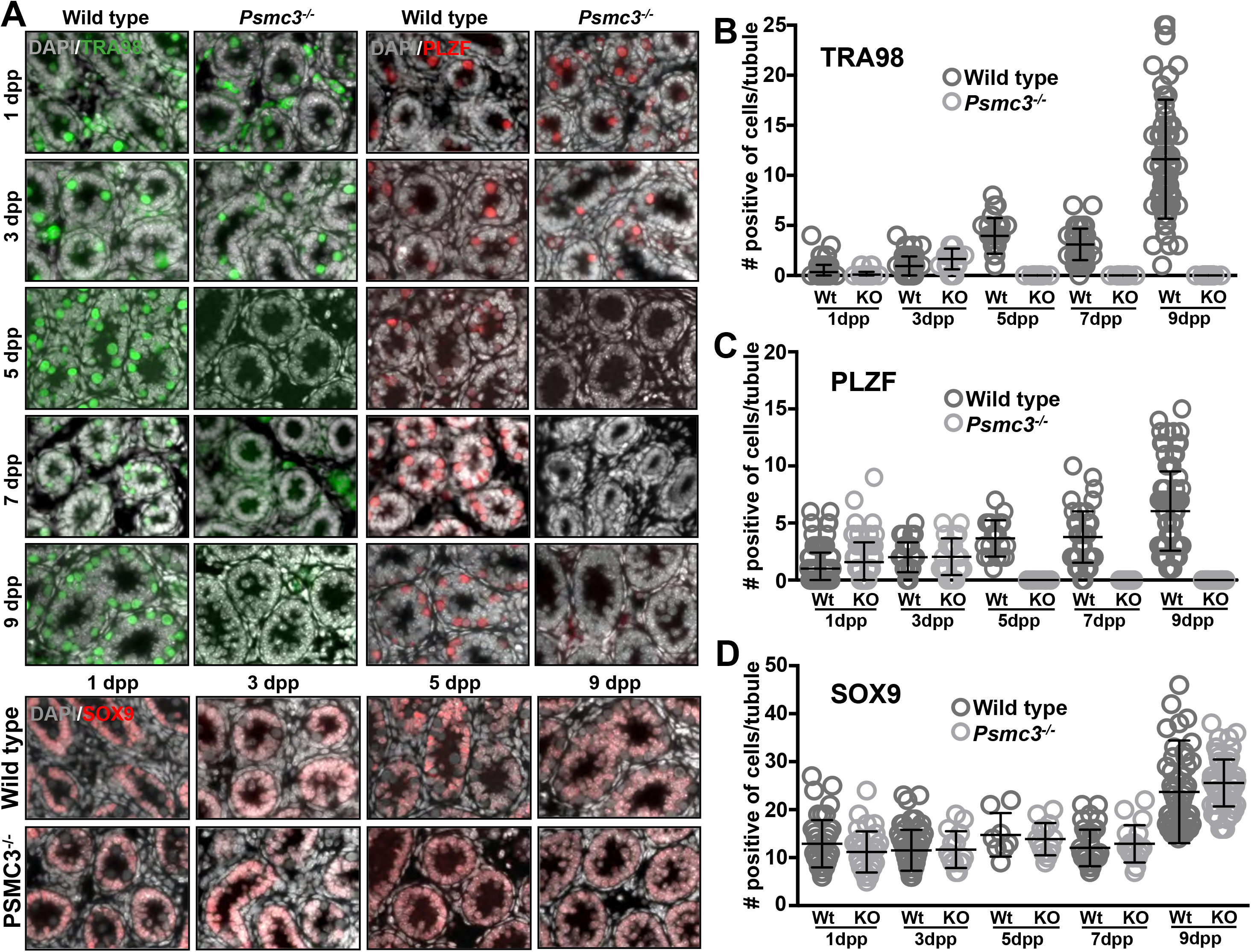
Testis cord differentiation defects in *Psmc3^−/−^* mice. **(A)** Histological sections of wild type and *Ddx4*-Psmc3^*−/−*^ testis cord from 1-9 days old mice immunostained with TRA98 antibodies (marking germ cells), PLZF antibodies (marking undifferentiated spermatogonia) and Sox9 (marking Sertoli cells). Quantitation of cells in A immunostained with TRA98 **(B)**, PLZF **(C),** and Sox9 (**D**).

Notably, compared to wild type (5dpp PLZF, 3.7 ± 1.6, n=20; TRA98, 3.9 ± 1.8, n=26. 7dpp PLZF, 3.8 ± 2.3, n=45; TRA98, 3.1 ± 1.5, n=48. 9dpp PLZF, 6.0 ± 3.5, n=86; TRA98, 11.62 ± 5.9, n=86) a significant reduction of PLZF and TRA98 positive cells in Psmc3^*−/−*^ mutants were observed at 5ddp (PLZF, 0.0 ± 0.0, n=20, P<0.0001, t test; TRA98, 0.0 ± 0.0, n=39, P<0.0001, t test) and 7dpp (PLZF, 0.0 ± 0.0, n=41, P<0.0001, t test; TRA98, 0.0 ± 0.0, n=41, P<0.0001, t test) and confirmed at 9dpp (PLZF, 0.0 ± 0.0, n=86, P<0.0001, t test; TRA98, 0.0 ± 0.0, n=94, P<0.0001, t test) testis (Fig. 4A-C). We note that the reduction in the number of positive cells for PLZF indicates that deletion of *Psmc3* affects undifferentiated stages of gamete development.

Since normal number of Sertoli cells revealed by immunostaining with SOX9 can be observed in testis of this mutant (3dpp wild type, 11.5 ± 4.3, n=58 and mutant 11.7 ± 3.8, n=15, P=0.9, t test. 5dpp wild type, 14.8 ± 4.5, n=15 and mutant 13.9 ± 3.5, n=14, P=0.6, t test. 7dpp wild type 12.0 ± 3.8, n=37 and mutant 12.9 ± 3.9, n=18, P=0.44, t test. 9dpp wild type 23.7 ± 10.7, n=64 and mutant 25.6 ± 4.9, n=75, P=0.18, t test) (Fig. 4A and D), we conclude that deletion of *Psmc3* by *Ddx4*-Cre affects germ cells at the undifferentiated stage of spermatogonia.

## Discussion

PSMC3 has been associated with a number of different cell functions, and it is highly expressed in testis. Nonetheless, PSMC3 function in gametogenesis is poorly understood. In this work, we took to task the function of PSMC3 in mouse gamete development. We generated and analyzed the phenotype of gonad-specific conditional *Psmc3* knockout mice. We observed that knocking out *Psmc3* in mouse spermatocytes results in severe male and female gonad developmental defect. Arrest of gametogenesis occurs at early pre-meiotic stages, revealed by a massive loss of undifferentiated spermatogonia, and apparently as a result of abnormal spermatogonia niche establishment and/or maintenance. Our results are in agreement with previous works showing that mutants affecting *Rpt5a* Arabidopsis (ortholog of *Psmc3*), result in severe male gametophyte defects, with pollen development arrested before cells enter meiosis at the second pollen mitosis stage. This correlates with absence of the proteasome-dependent cyclin A3 degradation and argues that gametophyte development may require proteasome function through RPR5A [13].

PSMC3 has also been associated to proteasome-independent functions. Indeed, PSMC3/TBPinteracts with HOP2/TBPIP [25], a central player in the meiotic recombination pathway. Deletion of HOP2 in mouse results in male and female gamete developmental defects, with impairment in double-strand break repair and homologous chromosome associations [28]. Because HOP2/TBPIP is a strong interactor of PSMC3, we reason that deletion of *Psmc3*, which may affect HOP2 integrity or activity, may result in meiotic defect. To test this, we analyzed *Spo11-Psmc3^−/−^* mouse testis, in which conditional deletion of *Psmc3* is predicted to occur at the onset of primary spermatocytes, after normal mitotic divisions and developing gametes have entered meiosis. Evaluated by the normal development of *Spo11-Psmc3^−/−^* testis, we conclude that PSMC3 is dispensable for normal meiotic progression, including double-stand break repair and homologous chromosome interactions.

In conclusion, our work defines a fundamental role of PSMC3 in spermatogenesis during early spermatogonia development. Future work, should address the mechanism of such function, either related or independent of PSMC3 participation in the proteasome.

